# Comparison of Efficiency and Specificity of CRISPR-Associated (Cas) Nucleases in Plants: An Expanded Toolkit for Precision Genome Engineering

**DOI:** 10.1101/422766

**Authors:** Oleg Raitskin, Christian Schudoma, Anthony West, Nicola J. Patron

## Abstract

Molecular tools adapted from bacterial CRISPR (Clustered Regulatory Interspaced Short Palindromic Repeats) systems for adaptive immunity have become widely used for plant genome engineering, both to investigate gene functions and to engineer desirable traits. A number of different Cas (CRISPR-associated) nucleases are now used but, as most studies performed to date have engineered different targets using a variety of plant species and molecular tools, it has been difficult to draw conclusions about the comparative performance of different nucleases. Due to the time and effort required to regenerate engineered plants, efficiency is critical. In addition, there have been several reports of mutations at sequences with less than perfect identity to the target. While in some plant species it is possible to remove these so-called ‘off-targets’ by backcrossing to a parental line, the specificity of genome engineering tools is important when targeting specific members of closely-related gene families, especially when recent paralogues are co-located in the genome and unlikely to segregate. Specificity is also important for species that take years to reach sexual maturity or that are clonally propagated. Here, we directly compare the efficiency and specificity of Cas nucleases from different bacterial species together with engineered variants of Cas9. We find that the nucleotide content correlates with efficiency and that Cas9 from Staphylococcus aureus is comparatively most efficient at inducing mutations. We also demonstrate that ‘high-fidelity’ variants of Cas9 can reduce off-target mutations in plants. We present these molecular tools as standardised DNA parts to facilitate their re-use.

## Introduction

Components of bacterial CRISPR (Clustered Regulatory Interspaced Short Palindromic Repeats) systems for adaptive immunity have been repurposed for engineering the genomes of eukaryotic organisms (1–3). These molecular tools have been rapidly and successfully applied in many organisms, including plants (for recent reviews see Gao, 2018; Ricroch *et al.*, 2017; Yin *et al.*, 2017), primarily due to the ease at which they can be programmed to recognise new genomic targets. The majority of plant genome engineering studies have utilised Cas9 (CRISPR Associated Protein 9) from *Streptococcus pyogenes*, a monomeric nuclease found in the Type II CRISPR system of that species. The Cas9 protein can be directed to selected genomic targets by an engineered RNA moiety known as the single guide RNA (sgRNA) (1,2). One or more sgRNAs can be programmed to recognise new genetic targets by recoding the ~19 base pairs at the 5’ end of the molecule, known as the spacer (7). The sgRNA forms a ribonuclease complex with Cas9 enabling it to scan DNA, pausing when it encounters a cognate sequence known as the protospacer adjacent motif (PAM) (8). On recognition of a cognate PAM, the ribonuclease complex probes the adjacent sequence to determine identity to the spacer region. If complementary, the spacer region of the sgRNA forms a Watson-Crick base-pair with the target DNA forcing the Cas9 protein to undergo a conformational change that enables the nuclease domains to cleave each of the DNA strands (8). The most common application in plants has been targeted mutagenesis achieved following transgenic expression of sgRNA-guided Cas9 to introduce double strands breaks (DSBs) at selected genomic loci. These induced breaks are predominantly repaired by the endogenous mechanism of non-homologous end-joining (NHEJ), which sometimes introduces errors (9,10). Since a perfect repair will continue to be recognised by the Cas9/sgRNA complex, and therefore cut again, constitutive expression eventually results in mutations, typically small insertions or deletions at the target. In many transgenic events, these mutations occur sufficiently early in the development and regeneration of the transgenic plant that all cells of the plant contain the same mutant genotype. A smaller number of studies have successfully leveraged the induction of targeted DSBs to increase the efficiency of targeted integration via the mechanism of homology directed repair (HDR), enabling transgenes to be inserted at a precise locus or for genomic DNA sequences to specifically recoded (for example, Baltes *et al.*, 2014; Begemann *et al.*, 2017; Cermak *et al.*, 2017; Čermák *et al.*, 2015; Gil-Humanes *et al.*, 2017; Li *et al.*, 2013).

Additional Cas proteins from other bacterial species have been adapted for genome engineering in eukaryotes and have also been applied to plants. These include Cas9 from the *Staphylococcus aureus* Type II CRISPR system (17–19) and Cas12a (previously Cpf1) from the Type V CRISPR systems found in *Francisella novicida, Acidaminococcus sp. and Lachnospiraceae bacterium* (3,14,20,21). Cas proteins from different species generally show preferences for different PAMs and, therefore, these additional Cas proteins have increased the number of genomic sites that can be targeted for engineering. While SpCas9 most efficiently cleaves DNA adjacent to NGG PAMs, SaCas9 is reported to show preference for NNGRRT (Friedland *et al.*, 2015; Kleinstiver, *et al.*, 2015a; Xie *et al.*, 2018). Cas12a-RNA complexes efficiently cleave target DNA preceded by a short T-rich PAM, with data suggesting a preference for TTTV (3,25). In addition, engineered versions of Cas9 and Cas12a with mutations in their PAM recognition domains have further expanded the repertoire of target sites to include sequences adjacent to NGAG and NGCG PAMS (with variants of SpCas *et al.*, 2015b) and NNNRRT PAMs (with variants of SaCas9 (Kleinstiver, *et al.*, 2015a)).

Cas-mediated genome engineering in plants has often utilised established transformation methods (e.g. *Agrobacterium*-mediated) to deliver DNA molecules encoding expression cassettes for the Cas9 protein, one or more guide RNAs and, typically, a plant selectable marker cassette, to the plant cell aiming for integration of all components at a single genetic locus. To achieve this, several systems have been developed for the assembly of multigene constructs, many of which employ Type IIS restriction cloning, also known as Golden Gate assembly (11, 27–31). Following the recovery of transgenic plants, most studies report the ‘efficiency’ of targeted mutagenesis as the percentage of transgenic plants in which mutations are found at the intended target. Most published studies have utilised different genetic targets, construct designs and regulatory elements making it difficult to draw conclusions about the comparative efficiencies and specificities of each nuclease. A few recent studies have attempted to compare Cas9 and Cas12a nucleases in similar experimental conditions (e.g. Lee *et al.*, 2018). However, they have necessarily used different targets with each type of nuclease, which are known to exert significant influence on efficiency (33).

Another area of interest has been the specificity of Cas nucleases. With careful bioinformatic analysis it is often possible to identify target sites in the genomic region of interest that are unique within the genome, thereby reducing the likelihood of mutations being induced at alternative loci. However, a number of different studies, including several in plants, have reported so-called “off-target” mutations at sites in the genome with less than perfect identity to the spacer (34–36). In some cases, small numbers of off-target mutations are of no concern, especially as mutations are known to occur during the process of tissue culture and regeneration (37,38). Further, many species can be back-crossed to the parent to remove unwanted mutations. However, when the aim is to induce mutations in specific members of a gene family, specificity is desirable, especially for recent paralogues that may be in close proximity and therefore impossible to segregate. In addition, the introduction of additional unwanted mutations in lineages that are not typically sexually propagated, such as cultivars of *Solanum tuberosum* (potato), is particularly undesirable.

To increase the specificity of Cas9 proteins for their target, researchers have engineered the nuclease and PAM-recognition domains (39–41). The resulting proteins were initially tested in mammalian cell cultures and reported to maintain similar levels of efficiency to the wild-type protein at sequences with an exact match to the spacer, but limited activity at sequences with less than perfect identity to the spacer. Here we report the outcomes of experiments that compare the efficiency and specificity of multiple wild type and engineered variants of Cas nucleases. All constructs used exactly the same regulatory elements and are assembled into identical vectors for delivery. Importantly, the efficiency and specificity of each nuclease are initially compared at the same target and, by quantifying the frequency of mutagenesis induced during transient expression, we avoid the influence of transgene insertion location on transgene expression. We also present an analysis of all potential targets as well as their off-targets in the coding exons of Arabidopsis and test our tools at a larger number of targets to identify factors that influence efficiency.

## Methods

### Analysis of targets in Arabidopsis coding sequences

Coding sequences were extracted from the *Arabidopsis thaliana* TAIR10 annotated whole chromosome datasets (ftp://ftp.arabidopsis.org/home/tair/Sequences/whole_chromosomes/ https://www.arabidopsis.org/download_files/Genes/TAIR10_genome_release/TAIR10_gff3/TAIR10_GFF3_genes.gff). Jellyfish (v.2.2.6) (42) was combined with a script to extract all possible legitimate candidate target sequences for each Cas nuclease [N_20_NGG (SpCas9), N_21_-NNGGGT (SaCas9), N_20_-NGAG (SpCas9-VQR), N_20_-NGCG (SpCas9-VRER), TTTV-N_23_ (Lb/AsCas12a)]. Potential off-targets (target sequences that are either identical to another target or differ by, at most, one base pair in the spacer region) were detected by mapping all identified targets against the TAIR10 genomic sequences using bbmap (v.38.06) with the following parameters (k=8, minid=0.75, ambig=all, mappedonly=t, secondary=t, sssr=0.75, ssao=t, saa=f, mdtag=t, nhtag=t, xmtag=t, amtag=t, nmtag=t, xstag=t, indelfilter=0, subfilter=4, editfilter=4). Alignments were filtered by Hamming distance allowing at most one mismatch in the spacer (plus mismatches at the ambiguous positions of the respective PAM sequences). Any potential off-targets located in coding sequences were identified with Bedtools intersect (v. 2.26.0; Quinlan and Hall, 2010) requiring an overlap equal to the length of the target including the PAM. All identified targets are provided in Supplementary File 1. All scripts and a snakemake pipeline (44) containing the whole workflow are available at https://github.com/EI-CoreBioinformatics/CRISPRanto.

### Selection of targets and off-targets for assessment of targeted mutagenesis

To remove any potential variability that might be associated with gene expression, we selected targets/off-target pairs located in genes expressed in leaves. To do this, we compared our candidate target/off-target pairs to gene expression data from the Expression Atlas (https://www.ebi.ac.uk/gxa/home, experiment E-GEOD-38612), retaining those targets within genes expressed in leaves. We then selected candidate target/off-target pairs that differed within the first 3 bp distal to the PAM. Finally, we either selected target/off-target pairs in which the expected cleavage site overlapped with the recognition site for a Type II restriction endonuclease, or searched 1000 bp of sequence each side of the target/off-target pairs to identify the presence of an additional target, identical in both sequences) that would enable the creation of a deletion when a second sgRNA recognising this target was delivered.

### Assembly of constructs

Constructs were assembled using the Plant Modular Cloning (MoClo) plasmid toolkit (45), a gift from Sylvestre Marrillonet (Addgene Kit # 1000000044). New Level 0 parts were made according to the standards described in (46). All Cas9 proteins were fused at the C-terminus to yellow fluorescent protein (YFP) and nuclear localisation (NLS) KKRKVKKRKVKKRV) tags. Cas12a proteins were fused to the same C-terminal NLS tag, followed by a 3xHA tag. Level 0 parts were assembled into transcriptional units in Level 1 acceptor plasmids in a one-step digestion-ligation reaction, except for sgRNAs, in which a U6 promoter L0 part was assembled into Level 1 acceptor plasmid together with a PCR amplicon of the complete sgRNA (spacer and scaffold), as described in (35). Subsequently, Level 1 transcriptional units were assembled into the Level 2 acceptor plasmid pICSL4723 (Addgene #86172). The digestion-ligation reactions were set up either manually or at nanoscale using laboratory automation: For manual assembly, 15 fmol of each DNA part was combined with 7.5 fmol of the acceptor plasmid in a final volume of 5 μL dH_2_0. This was combined with 5 μL of reaction mix (3 μL of dH_2_0, 1 μL of T4 DNA ligase buffer 10x (NEB, Ipswich, MA, USA), 0.5 μL of 1 mg/mL purified bovine serum albumin (1:20 dilution in dH_2_0 of BSA, Molecular Biology Grade 20 mg/mL, NEB), 0.25 μL of T4 DNA ligase at 400 U/μL (NEB) and 0.25 μL of *BsaI* or *BpiI* restriction enzyme at 10 U/μL (ThermoFisher, Waltham, MA, USA)) and incubated in a thermocycler for 26 cycles of 37°C for three minutes followed by 16°C for four minutes and a final incubation at 37°C followed for 5 minutes and 80°C for five minutes. A 2 μL aliquot of each reaction was transformed into 20 μL electrocompetent cells and plated on selective LB-agar plates. Automated reactions were scaled down to a final reaction volume of 1 μ using the Echo 550 liquid handler (Labcyte Ltd. San Jose, SA, USA), transformed into 2 μL XL10-Gold^®^ Ultracompetent Cells (Agilent Technologies, Santa Clara, CA, USA) and plated onto eight-well selective LB-agar plates on a Hamilton^®^ STARplus platform. The sequences of assembled plasmids were verified by complete sequencing using 150 base pair paired-end reads on an Illumina MiSeq platform. Libraries were prepared using the Nextera XT DNA Library Prep Kit (Illumina, San Diego, CA, USA) with a modified 2 μL total volume protocol using a one in 25 dilution of components. A complete list of all 132 plasmids, comprising Level 0 DNA parts, Level 1 transcriptional units and Level 2 constructs, used in this study is given in Supplementary File 2. Samples, together with complete annotated sequence files for the 128 new plasmids generated for this study have been deposited at the AddGene plasmid repository.

### Transient expression in protoplasts

For each experiment, a sufficient number of protoplasts were prepared to enable the delivery of four replicates of all constructs to be compared. Protoplasts were prepared from the leaf tissues of *Nicotiana benthamiana* or *A. thaliana* as previously described (47). Protoplasts were quantified and divided into aliquots of 200 μL in transfection buffer (0.4M mannitol, 15mM MgCl_2_, 4 mM MES, pH 5.6), each containing approximately 1 x 10^4^/ml intact protoplasts, such that four separate aliquots of protoplasts from the same preparation were available for each of the plasmids to be compared. Plasmid DNA for delivery to protoplasts was prepared using the Plasmid Plus Midi kit (Qiagen, Hilden, Germany) with a modified protocol incorporating three additional wash steps prior to elution from the column. Freshly made PEG (2 g of PEG (Mn 4000 (Sigma, 81240)) in 2 mL of 500 mM mannitol and 0.5 mL of 1M CaCl_2_) was mixed with10 μg of purified DNA and added to each aliquot of protoplasts. Subsequently, protoplasts were washed and resuspended in 300 μL of washing buffer (154 mM NaCl, 125 mM, CaCl_2_, 5 mM KCl, 2 mM MES; pH5.6) and incubated for 24 hours at 24°C in an illuminated incubator with light intensity of approximately 70 μmol/m^2^/s. Transformation efficiency was estimated by quantification of protoplasts in which YFP fluorescence was visible in the nuclei using an inverted fluorescence microscope (Zeiss Axio Observer Z1 or ThermoFisher Evos).

### Detection and quantitation of targeted mutagenesis

DNA was extracted from the protoplasts using a cetyltrimethylammonium bromide (CTAB) extraction protocol: pellets of protoplasts were resuspended in 100 μL of extraction buffer (0.2M Tris-HCl, pH7.5; 0.05M EDTA; 2M NaCl; 2% CTAB, pH7.4) and incubated at 65°C for 1 hour prior to addition of 45 μL chloroform. Following centrifugation, the upper aqueous phase was precipitated with an equal volume of isopropanol. DNA pellets were washed with 70% w/v ethanol, dried and resuspended in sterile distilled water with 5 μg/μL RNAse A. Each target was amplified using a pair of primers, specific to the locus of interest (Supplementary File 3). PCR reactions were performed using 70 ng DNA and Q5 High-Fidelity DNA Polymerase (NEB) according to the instructions provided by the manufacturer. Mutations at the targets were identified by either Illumina or Sanger sequencing. For preparation for Illumina sequencing, amplicons were purified using Agencourt AMPure XP (Beckman Coulter) and indexed using the Nextera XT Library Preparation kit (Illumina) according to the manufacturer’s instructions. Reactions were analysed by microfluidic gel fractionation (LabChip GXII, Perkin Elmer) and pooled. Primers were removed by fractionation (BluePippin, SAGE Science). The concentration and quality of DNA was analysed by QUBIT (ThermoFisher) and qPCR (Kapa Library Quantification Kit, Illumina). phiX Sequencing Control V3 (Illumina) was added to final concentration of 1.75 pM. Sequencing was performed on an Illumina MiSeq using the MiSeq Reagent Kit v2 Micro. Adapter sequences were removed and quality trimmed using bbduk (v.37.24) (https://jgi.doe.gov/data-and-tools/bbtools/bb-tools-user-guide/bbduk-guide/) (parameters ktrim=r k=21 mink=11 hdist=2 qtrim=lr trimq=3 maq=10 ftr=250). Quantification of mutations at the target was performed using CRISPRESSO (http://crispresso.rocks/) (48) and automated with a custom script (available at https://github.com/EI-CoreBioinformatics/CRISPRanto). The quantity of mutations in each sample was then normalised to the quantified transfection efficiency.

For analysis by Sanger sequencing, amplicons were purified (Qiaquick PCR purification, Qiagen) and incubated with a restriction enzyme for which the recognition sequence overlapped the expected cleavage site (see selection of targets) prior to reamplification. This reduced the amount of wild-type sequence in the sample, enabling detection of low-abundance amplicons. These amplicons were sequenced directly (Eurofins) and evidence of mutagenesis was conferred by the presence of multiple peaks, representing the different DSB-repair events across the population of cells, visible after the expected cut site (see Supplementary File 4). These chromatogram signals were analysed using the ICE software that determines rates of CRISPR-Cas9 editing at a specific, sgRNA directed genomic location within a cell population (Synthego, https://ice.synthego.com/).

## Results

### Codon optimisation and sgRNA structure have minimal effects on the efficiency of targeted mutagenesis in plants

Prior to comparison with other Cas nucleases, we first assessed several variables for RNA-guided Cas9. We assessed the effect of codon-optimisation, comparing human (SpCas-h) versus plant (SpCas9-p), as well as variations in the single guide RNA sequence, comparing two sgRNAs: sgRNA and an sgRNA with extended step-loops (sgRNA-ES; Chen *et al.*, 2013).

We also compared the use of a previously extended endogenous terminator for sgRNA expression cassettes (50,51). In these and subsequent experiments, all constructs were assembled similarly using identical regulatory sequences (Fig 1A). All sgRNAs contained a spacer to direct Cas9 to a target in the phytoene desaturate gene of *N. benthamiana* (*NbPDS1*). Following a quantitative assessment of the frequency of mutagenesis by Illumina sequencing, we found no significant differences in the number of sequencing reads with mutations at the target (Supplementary File 5), indicating that neither human codon optimisation, the shorter stems found in the original sgRNA, or the minimal terminator significantly impaired the efficiency of mutagenesis in plants. In all subsequent experiments, we used SpCas9-h together with its original sgRNA as we have had previous success with these sequence in other species (35).

**Figure 1.**
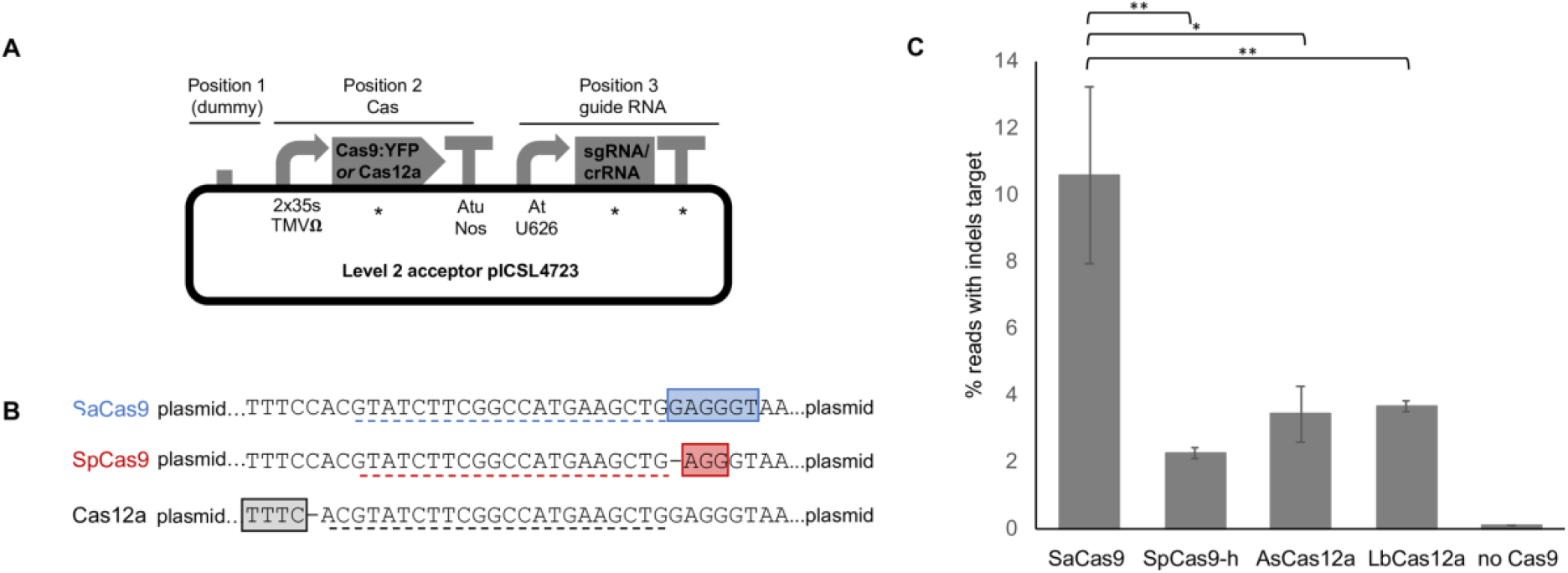
Comparison of Cas9 and Cas12 nucleases at a single target. (A) All constructs were assembled in the same backbone, using the same regulatory elements. Asterisks (*) represent elements compared in this study. The ‘dummy’ consists of 15 random nucleotides and enables the Cas9 and sgRNA expression cassettes to be reassembled with, e.g. a selectable marker cassette, in Position 1. (B) An identical target sequence was used to test all Cas proteins. The target was flanked by the preferred cognate protospacer adjacent motif (PAM) for SpCas9 (red box), SaCas9 (blue box) or Cas12a (black box). The spacer sequence used in each sgRNA (Cas9) or crRNA (Cas12a) are indicated by dotted lines. (C) Under identical experimental conditions, SaCas9 induced more mutations at the target than SpCas9, AsCas12a or LbCas12a. Error bars = standard error of the mean; n=4. Post hoc comparisons using Tukey’s Honest Significant Difference test indicted significant differences: * = p<0.05, ** = p<0.01.

### Cas proteins from different bacterial species show varied efficiencies of targeted mutagenesis at the same target in identical experimental conditions

To enable a direct comparison of the ability of four Cas proteins, SaCas9, SpCas9, AsCas12 and LbCas12a to induce targeted mutations, without the confounding influence of varied efficiency across targets, we designed three variants of a synthetic target such that the same recognition sequence was adjacent to the preferred PAM for each protein (Fig 1B). We then co-delivered the plasmid with the cognate PAM to protoplasts together with plasmid DNA encoding each of the four nucleases and an appropriate sgRNA (Cas9) or crRNA (Cas12a) with a spacer to the synthetic target. We observed significantly more mutations at the target with SaCas9 than with SpCas9-h, AsCas12 or LbCas12a (Fig 1C).

### Efficiency of mutagenesis correlates to GC content of the spacer

Analysis of the Arabidopsis genome for potential target sequences identified 3,853,090 potential targets for SpCas9, SaCas9 or Lb/AsCas12a in coding exons (Table1). Of the 2,695,798 targets recognised by SpCas9, 61,739 (2.29 %) have at least one potential off-target (identical or differing at only a single base) also in a coding exon, 7,201 of which differed by a single base in the first three positions. The total number of targets is expectedly lower for SaCas9 and Lb/AsCas12a, which recognise longer PAMs, (Table 1). We filtered these targets for those in genes previously shown to be expressed in leaves (see methods) and selected sequences for functional analysis in Arabidopsis leaf-derived protoplasts.

**Table 1.**
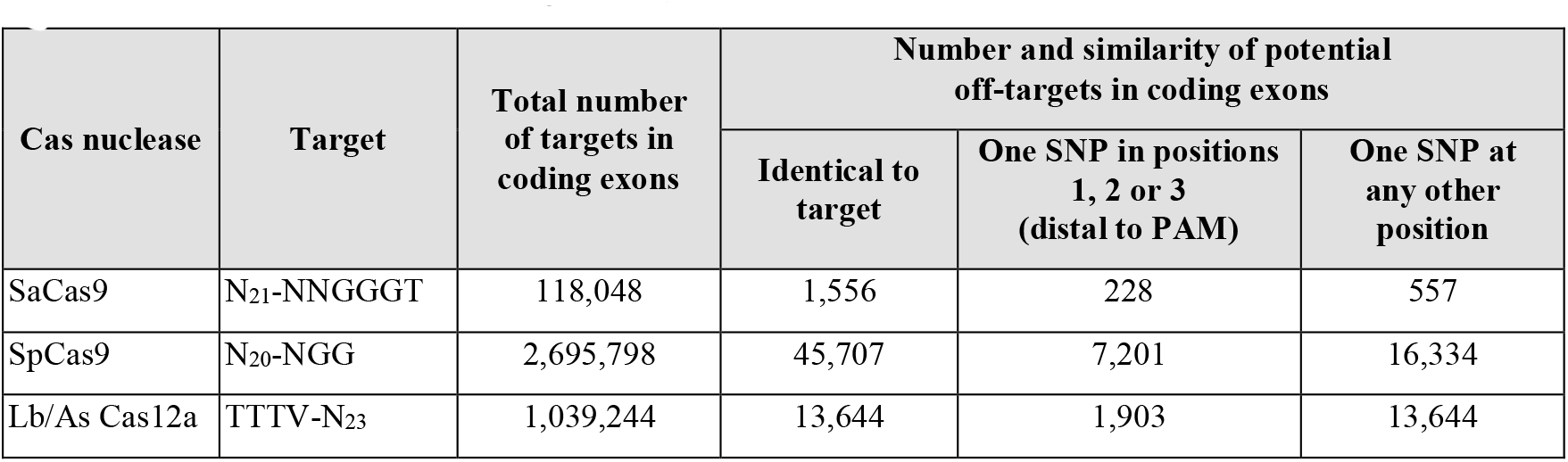
Numbers of Cas9 and Cas12 targets and potential off-targets in Arabidopsis coding exons.

We were able to detect evidence of induced mutations at 14 out of 33 candidate targets for SpCas9. We were able to detect evidence of induced mutations at seven out of nine candidate targets for SaCas9. The sample set for SaCas9 is too small for meaningful analysis, however, analysis of nucleotide composition of the 33 SpCas9 targets found that the %GC content of spacers used where mutagenesis was detected was significantly greater than those for which no activity was detected (single-tailed T-test, P<0.05) (Fig 2). The data indicates that GC content of spacers should, for maximal efficiency, be greater that 40%. In contrast, we found no differences in the GC content within the seed region.

**Fig 2.**
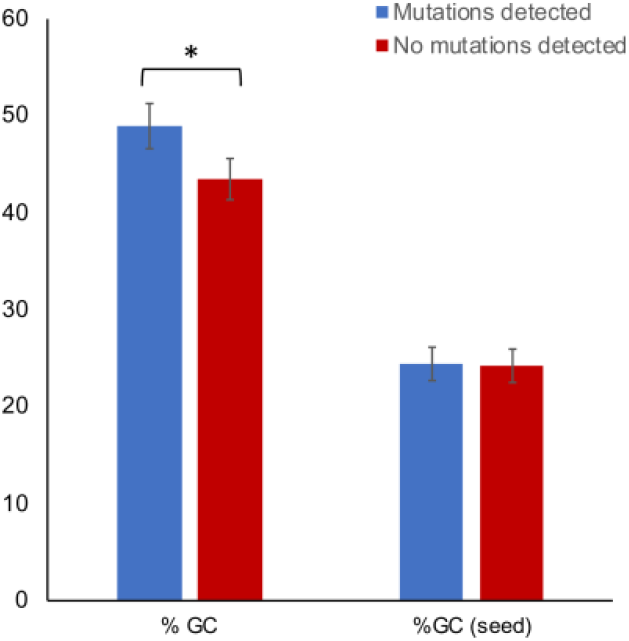
Nucleotide content analysis of 33 spacer sequences. All spacers have cognate targets in the coding sequences of leaf-expressed genes tested in sgRNAs with SpCas9-h. Each spacer was incorporated into a sgRNA and tested in Arabidopsis protoplasts. Blue bars indicate mutations were detected at target; red bars indicate no mutations detected at target. Seed = six base pairs adjacent to PAM. *= p-value 0.044591. The result is significant at p <0.05.

### ‘High-fidelity’ variants of Cas9 show reduced efficiency at targets with less than perfect identity to the spacer

To directly compare the efficiency and specificity, *in planta*, of five variants of SpCas9 nucleases, we used a similar experimental process as reported by (Kleinstiver, *et al.*, 2016). We designed a set of five sgRNAs each with a mutation in a different base of a spacer designed to target the *NbPDS* gene (Fig 3A). An sgRNA with an exactly matching spacer, or one of the five variants was delivered to plants cells in combination with each of the five variants of SpCas9 and the number of targeted mutations was quantified using Illumina sequencing. While the number of mutations induced by wild type Cas9 was significantly reduced when the spacer contained a mutation in the region close to the PAM, the presence of a mismatch between the spacer and target in the distal region had minimal effects (Fig 3B). In contrast, the frequency of mutations induced by the variants eCas9 1.0 and 1.2 was significantly reduced by a mismatch in any region of the sgRNA (Fig 3B). This experiment, although quantitative, was difficult to scale across a larger number of targets. To compare the efficiency of mutagenesis across a larger number of targets and to observe the performance of spacers at non-identical endogenous targets, we conducted an analysis of the Arabidopsis genome to identify pairs of targets that differed by a single base pair. For each target at which mutagenesis was detected using SpCas9-h or SaCas9, the same sgRNA were also tested with protein variants (eCas9 1.0, eCas9 1.1, eSaCas9 and the recently reported xCas9 3.7) described to have reduced activity at targets with a less than perfect match to the spacer. As well as analysis at the targets, we also analysed the identified off-target locus for evidence of mutagenesis (Table 2). Of eight targets at which SpCas9-h induced mutations at the target, mutations were also detected at a second target that differed by one nucleotide in the first three positions. Of these eight, eCas9 1.0 was only able to induce mutations at three targets but off-target activity was detected at two of these. Similarly, eCas9 1.1 was able to induce mutations at five targets but off-target activity was detected at four; xCas9 was able to induce mutations at three targets but off-target activity was detected at two of these (Table 2 and Supplementary File 4). In general, the signal of mutagenesis using the protein variants was less than for the equivalent wild-type protein sequence (Table 2 and Supplementary File 4).

**Fig 3.**
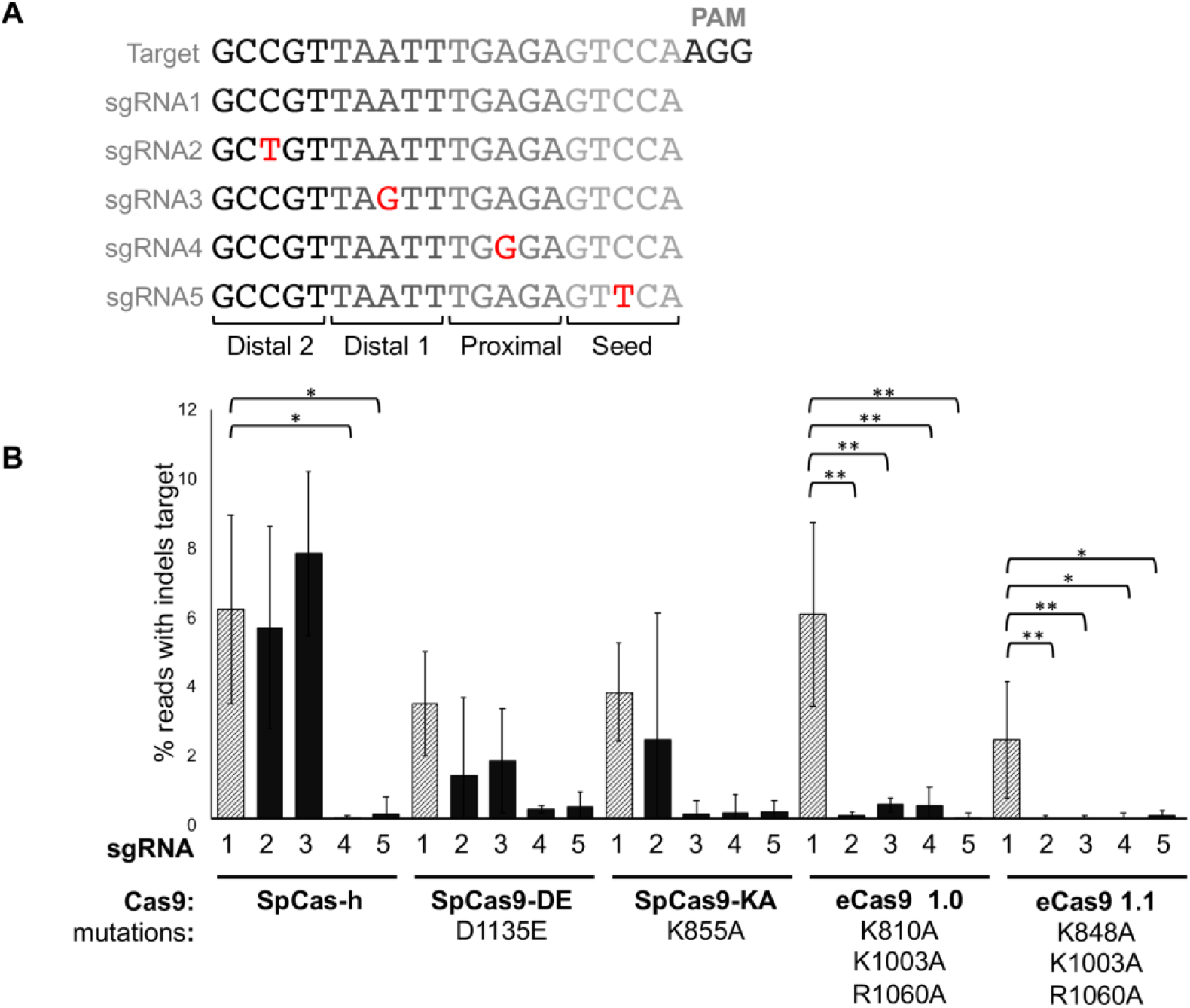
Assessment of Cas9 specificity at an identical target. **(A)** Schematic showing transversions in four regions of the spacer sequences of sgRNAs targeting *NbPDS1* **(B)** The efficiency of targeted mutagenesis is impacted by transversions in the spacer. Shaded bars represent a perfect match of the spacer to the target, black bars show sgRNAs with spacers with transversions relative to the target. Error bars = standard error of the mean; n=4. Post hoc comparisons using Tukey’s Honest Significant Difference test indicted significant differences: * = p<0.05, ** = p<0.01.

**Table 2.**
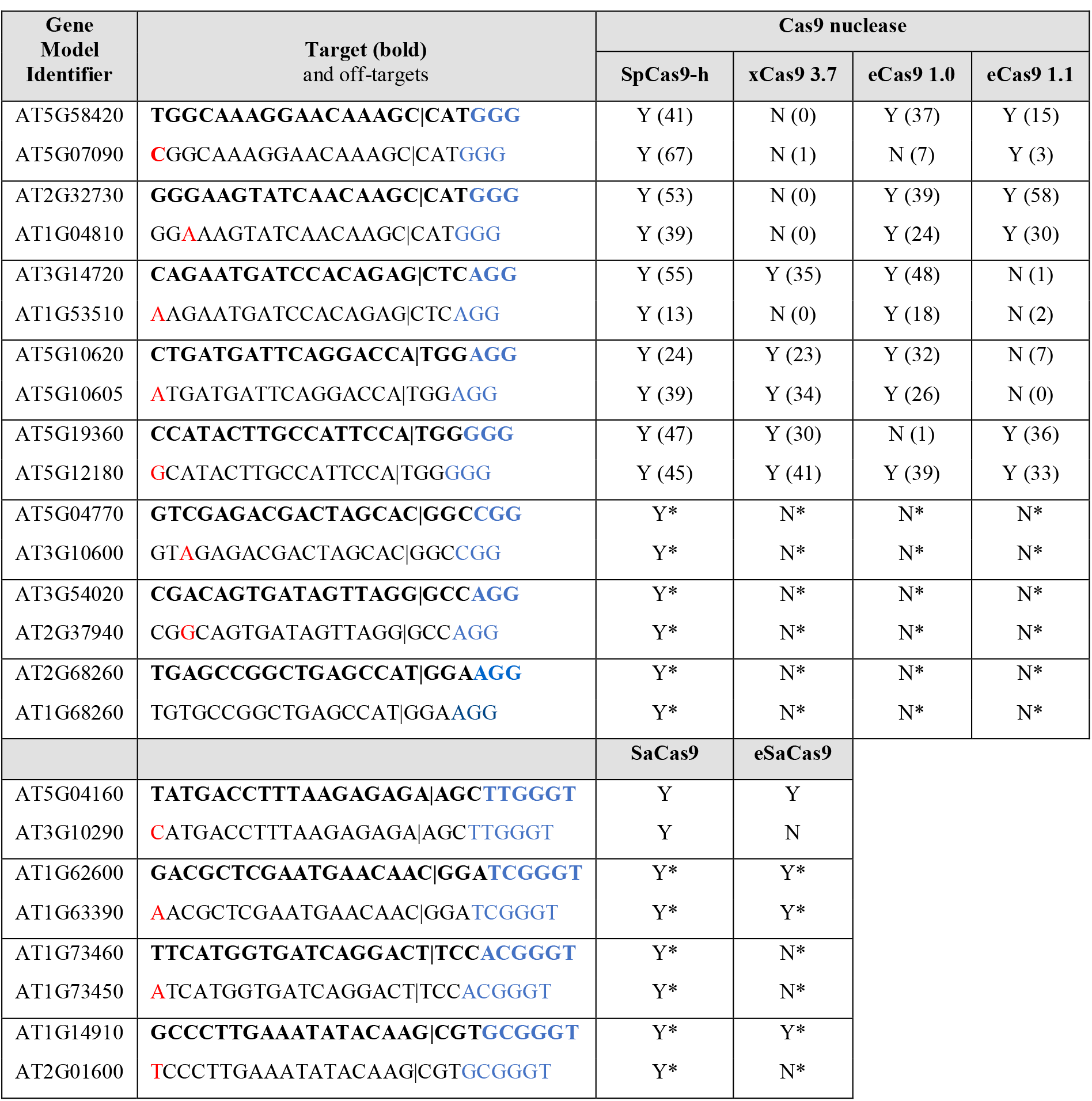
Assessment of the specificity of Cas9 variants at targets in Arabidopsis coding sequences. Differences between target (bold text) and off-target pairs are shown in red text. The Protospacer Adjacent Motif (PAM) is shown in blue text.Numbers in parentheses are scores from Inference of CRISPR Editing (ICE) software. “*” indicates assessment made by detection of deletion induced with a second sgRNA. “**|**” indicates expected point of cleavage.

## Discussion

Components of bacterial CRISPR/Cas systems have been applied to a wide variety of model and economically-important plant species including dicotyledonous fruit crops such as strawberries (52), tomatoes (53) and oranges (54) as well as monocotyledonous grain crops such as barley (35), wheat (55) and maize (56). In most cases, engineered plants are regenerated via somatic embryogenesis in tissue culture; a process that is both laborious and time-consuming. Consequently, researchers are interested in the efficiency of the molecular tools they deliver. Since the first reports of Cas9 from *S. pyogenes* being applied as tool for genome engineering, there has been much interest in novel properties associated with Cas nucleases from addition CRISPR systems (3,24). These include comparative efficiency and the potential for engineering at an increased number of genomic targets due to the recognition of additional PAMs. Because of this, Cas nucleases have often, necessarily, used different targets. Together with other differences introduced by different construct designs and experimental conditions, it is difficult to draw conclusions about relative efficiencies. Our experimental strategies allowed us to compare Much of the work on sequence determinants of spacers performed to date has been performed in mammalian cell cultures (59–63). These systems provide the advantage of being able to delivery libraries of sgRNAs targeting genes involved in a particular process followed by the selection of cells with the expected phenotype correlating to disruption of those genes to be selected (e.g. by staining and FACS). In an analysis of a number of datasets, including one of 4,000 sgRNA targeting 17 genes, Doench *et al.* (2016) concluded that the patterns of activity are complex, and are not likely to be apparent by examining smaller numbers of sgRNA:DNA interactions. Current methods of DNA delivery to plant cells include the preparation of separate strains of *Agrobacterium tumefaciens* for delivery to callus or tissues, or the direct delivery of plasmids, either to protoplasts or to tissues (using biolistic delivery). Delivery of a large-scale combined library of constructs using these methods would result in populations of cells with multiple induced mutations from which it would not be possible to identify or separate cells with individual genotypes. In our study, we delivered individual constructs to aliquots of protoplasts prepared from the same batch of leaf tissue. This allowed us to work with larger number of constructs, however, since cultures of protoplasts cannot be perpetuated (they reform cell walls and form masses of cells if allowed or induced to divide), it is not possible to select populations of cells beyond the first 24-48 hours. The advantages of this system, however, allowed us to compare the impact of a number of different components by allowing similar levels of expression across different experiments that are difficult to achieve with integrated transgenes, the expression of which is subject to copy number and the location of integration (64). Using this method, we attempted to induce mutations at over 45 targets in the Arabidopsis genome, using sgRNAs with spacers with both perfect and imperfect identity. This enabled us to compare the features of functional spacers (those that were able to induce targeted mutations) with non-functional spacer noting a correlation with GC content (Fig 2), broadly in agreement with that reported for mammals (59). We did not, however, note any determinants for sequence composition in the seed region or at specific bases (data not shown), likely because of the comparatively small size of our dataset. The correlation of efficiency with GC content may be related to nucleosome occupancy, previously reported to influence the ability of Cas9 to recognise its targets (65); nucleosome-depleted regions in Arabidopsis tend to have a higher GC content (66). As this negative correlation between nucleosome occupancy and GC content is not known to be common across all organisms, further studies in a wider range of plant species will need to be performed to determine if efficiency can be predicted by GC content alone As this negative correlation between nucleosome occupancy and GC content is not known to be common across all organisms, further studies in a wider range of plant species will need to be performed to determine if efficiency can be predicted by GC content alone.

Analysis of Arabidopsis coding sequences revealed that many targets for Cas-mediated genome engineering have one or more potential off-targets in other coding sequences that differ by just a single base (Table 1, S1). We provide experimental evidence that SpCas9 and SaCas9 are readily able to induce mutations at targets with less than perfect identity to the spacer (Table 2 and Fig 3). Different laboratories have engineered variants of SpCas9 and SaCas9, reportedly reducing their activity at targets to which the spacer does not have 100% identity (39–41, 67). To compare these proteins and determine their function in plants, we measured their efficiency and specificity across a number of targets (Table 2 and Fig 3). Unlike previous reports, we did not find that the use of precise perfectly matched guide sequences provided the same efficiency as the wild type protein (67) but that variants showed a reduction in efficiency at all targets. However, this was more substantial at sites where the target and spacer did not share 100% identity. In cases where specificity to a single target is critical e.g. targeting a specific member of a gene family, particularly when recent paralogues are located in tandem, these variants are likely to be useful.

This study utilised the ease of standardized modular cloning to create a suite of comparable constructs to enable direct comparison of multiple different tools for genome engineering. Type IIS mediated assembly methods have been widely utilised to facilitate the construction of the complex plasmids required for multiplexed genome editing (11, 29–31). In addition to standardising our experimental process, we are able to contribute an expanded toolkit of modular, reusable parts for plant genome engineering that will facilitate their application in new studies (Fig 4). The basic (Level 0) parts are flanked with inverted *BsaI* sites that will release parts with overhangs in the common genetic syntax for plants (46), making them amenable for reuse with a number of different assembly toolkits for plants.

**Fig 4.**
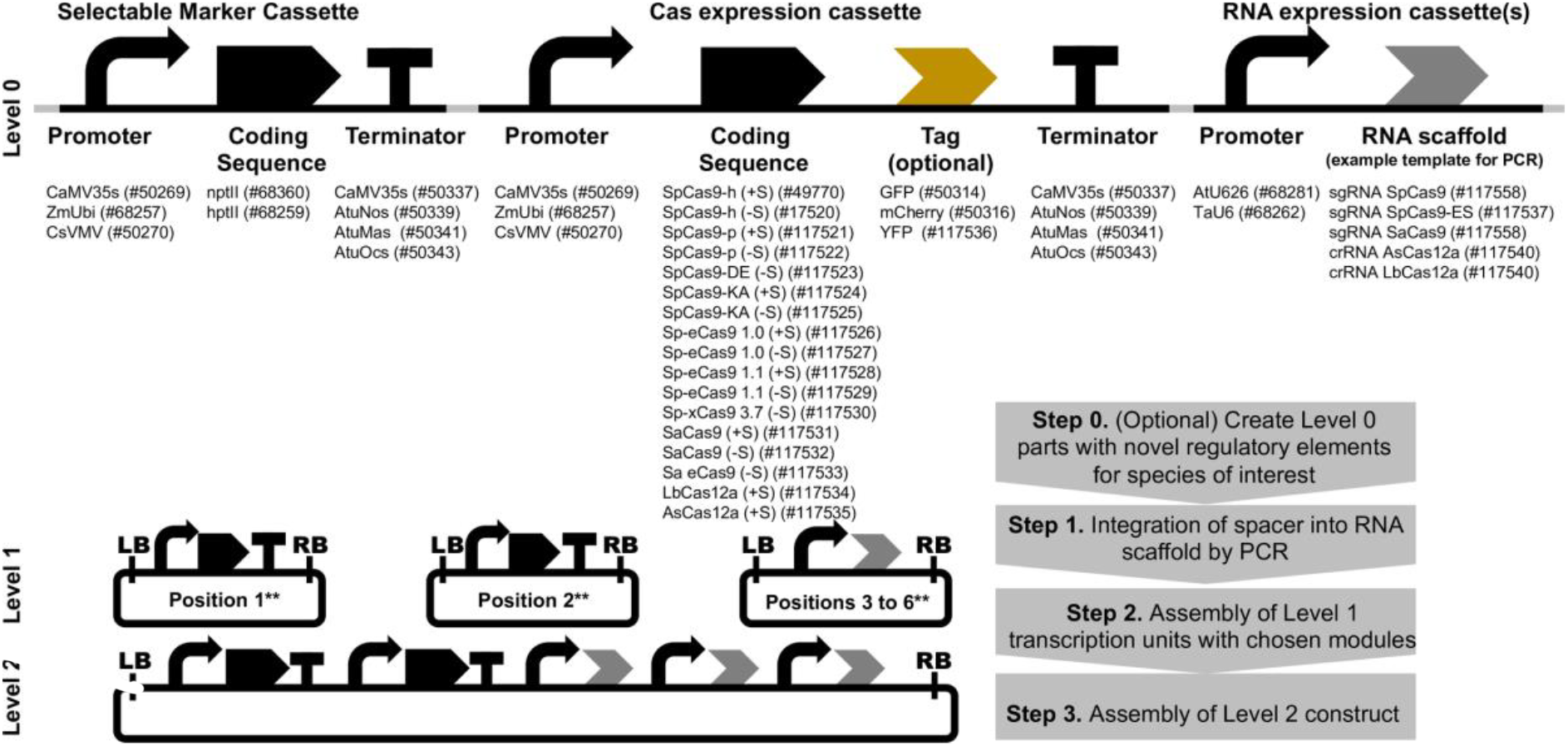
An expanded toolkit for Cas-mediated genome engineering in plants. Regulatory elements and coding sequences are cloned as Level 0 parts, enabling one-step assembly into transcriptional units mediated by *BsaI* (e.g. using the Plant MoClo Toolkit, Engler et al., 2014) and subsequently into multigene constructs. Numbers in parentheses represent catalogue number at Addgene. **For simplicity, all cassettes are shown assembled on the forward strand. However, the orientation of any cassette in the final construct can be altered by use of a reverse Level 1 acceptor. ‘+S’= with stop codon ‘-S’= no stop codon.

## Acknowledgements

This work was funded by the UK Biotechnological and Biological Sciences Research Council (BBSRC) and Engineering and Physics Research Council (EPSRC) Synthetic Biology for Growth program [OpenPlant Synthetic Biology Research Centre; BB/L014130/1]. Next-generation sequencing and library construction and automated DNA assembly was delivered via the BBSRC National Capability in Genomics at the Earlham Institute [BB/CCG1720/1]. CS was supported by BBSRC strategic funding, Core Capability Grant [BB/CCG1720/1, BBS/E/T/000PR9816].

## List of Supporting Information

**Supplementary File 1.** Details of genomic targets for Cas nucleases in *Arabidopsis thaliana* coding exons **Supplementary File 2.** Details of 135 plasmid constructs used in this study

**Supplementary File 3.** Table of oligonucleotide primers used to amplify genomic targets

**Supplementary File 4.** Chromatograms and gel images showing evidence of Cas-induced targeted mutations and deletions

**Supplementary File 5.** Effect of Cas9 codon-optimisation and sgRNA scaffold on the efficiency of targeted mutagenesis at *PDS1* in *Nicotiana benthamiana* protoplasts (h = human; p= plant; ES = extended stem; T= terminator). Error bars = standard error of the mean; n=4

